# TriPoly: a haplotype estimation approach for polyploids using sequencing data of related individuals

**DOI:** 10.1101/163162

**Authors:** Ehsan Motazedi, Dick de Ridder, Richard Finkers, Chris Maliepaard

**Affiliations:** Bioinformatics Group, Wageningen University and Research, The Netherlands; Wageningen UR Plant Breeding, Postbus 386, 6700AJ, Wageningen, The Netherlands

## Abstract

Knowledge of “haplotypes”, i.e. phased and ordered marker alleles on a chromosome, is essential to answer many questions in genetics and genomics. By generating short pieces of DNA sequence, high-throughput modern sequencing technologies make estimation of haplotypes possible for single individuals. In polyploids, however, haplotype estimation methods usually require deep coverage to achieve sufficient accuracy. This often renders sequencing-based approaches too costly to be applied to large populations needed in studies of Quantitative Trait Loci (QTL).

We propose a novel haplotype estimation method for polyploids, TriPoly, that combines sequencing data with Mendelian inheritance rules to infer haplotypes in parent-offspring trios. Using realistic simulations of short-read sequencing data for potato (*Solanum tuberosum*) and banana (*Musa acuminata*) trios, we show that TriPoly yields more accurate progeny haplotypes at low coverages compared to the existing methods that work on single individuals.

## 1 Introduction

Haplotypes are defined as sequences of consecutive nucleotides over a chromosome, which normally shares high similarity with *k* – 1 other chromosomes in diploid (*k* = 2) and polyploid (*k* > 2) organisms. These *k* homologous chromosomes can nevertheless have important differences in the form of nucleotide substitutions or insertions/deletions, leading to genotypic (and phenotypic) diversity within an outcrossing population, e.g. of the diploid (*k* = 2) human (*Homo sapiens*), tetraploid (*k* = 4) African clawed frog (*Xenopus laevis*) or tetraploid potato (*Solanum tuberosum*), or between inbred lines of autogamous species, e.g. hexaploid (*k* = 6) wheat (*Triticum aestivum*). The assignment of these variant forms, i.e. alleles, to the chromosomes is called *phasing* or *haplotyping*. In this context, phasing may also refer to the set of phased homologues, *H* = *{h*_1_, *h*_2_, *…, h*_*k*_*}* with *k* being the ploidy level and *h*_*i*_ (*i* = 1, *…, k*) being the haplotype corresponding to the *i*^th^ homologue.

As phasing is uninformative at genomic positions with identical nucleotides over all the homologous chromosomes, i.e. at the homozygous sites, haplotypes are usually defined as sequences of alleles at heterozygous sites over a chromosome. By this definition, 2^*n*^ haplotypes are theoretically possible for a region covering *n* bi-allelic Single Nucleotide Polymorphisms (SNPs), which is the most abundant form of genomic variation among individuals of the same species (Rafalski, 2002). However, often far fewer haplotypes actually occur in a study population.

While high-throughput genotyping assays such as SNP arrays can assist in efficient determination of unphased SNPs, direct determination of haplotypes is much more complicated due to high similarity of their nucleotide content, which usually requires the application of laborious and expensive techniques such as allele-specific PCR or chromosome microdissection (Michalatos-Beloin *et al.*, 1996; Doležel *et al.*, 2014).

However, unphased SNPs provide incomplete knowledge of an individual’s phenotype with respect to both gene expression and protein function, as both can be affected by the heterozygous variants being in *cis* or *trans* with other variants (Tewhey *et al.*, 2011). Besides, haplotypes can be used as multi-allelic markers offering more statistical power compared to single SNPs for genetic linkage and association studies (Simko *et al.*, 2004).

Several computational methods have been therefore proposed to indirectly infer the phasing from available genotype data, which can be divided into three main categories. Methods in the first category, such as *TetraOrigin* (Zheng *et al.*, 2016), aim to determine the most likely haplotypes using the segregation of marker alleles in a population taking into account the genetic distances between the marker loci. These methods start from unphased SNP data at positions far enough apart to be informative about linkage, and are especially useful with large populations (Garg *et al.*, 2016). Methods in the second category, such as *HapCut* (Bansal and Bafna, 2008), *HapCompass* (Aguiar and Istrail, 2013), *HapTree* (Berger *et al.*, 2014) and *SDhaP* (Das and Vikalo, 2015), exploit the fact that a sequence read containing at least two SNPs reveals the phasing of the homologue from which it has originated at the contained SNP sites. The aim of these methods is therefore to assign the reads of a single individual into *k* groups, corresponding to the homologues of a *k*-ploid, and to obtain the consensus sequence of the reads within each group to reconstruct the haplotypes over the sequenced region. Finally, methods in the third category are based on coalescence theory, trying to infer the haplotypes parsimoniously by minimising their total number in a population of unrelated (or only distantly related) individuals (Clark, 1990) or by applying data-augmentation to obtain a set of highly frequent haplotypes in the population compatible with the genotype data, as implemented in *SHEsisPlus* (Shen *et al.*, 2016).

Applied to polyploid species, all of these approaches have limitations in terms of applicable ploidy level (*k*), required marker density, sequencing depth and read length. For example, the TetraOrigin algorithm (the first category) is only applicable to bi-parental tetraploid populations (*k* = 4) with an obtained linkage map, and methods in the second category can fail to reconstruct haplotypes with high quality at low sequence depths as well as at ploidy levels higher than *k* = 4 (Motazedi *et al.*, 2017).

In case parent-progeny relations exist in a population, it is possible to improve the quality of haplotype estimation by combining the phasing information used in the first and second categories under a unified scheme. Such an approach is also of high practical importance, as with sequencing experiments becoming cheaper and more efficient, more often whole populations are sequenced rather than only genotyped at specific marker loci. An implementation of this unifying framework, called *PedMEC*, has recently been reported by Garg *et al.* (2016) for diploid *trios*, i.e. families with two parents and one offspring. Specifically, PedMEC extends the partial-phasing of sequence reads using their overlaps while penalising meiotic recombination events in each trio. However, the exact dynamic programming approach of Garg *et al.* (2016) rapidly becomes intractable for polyploids, i.e. with *k* > 2, as its complexity increases exponentially with an increase in the ploidy level (Section 2.3). Here we present a greedy algorithm, *TriPoly*, for phasing of the SNPs detected over a continuous genomic region in parent-offspring trios. Starting at the SNP site with the smallest genomic coordinate, TriPoly extends the phasing one SNP at a time keeping only the most likely extended phasings to be worked out in the subsequent extension step. In determining the likelihood of each extension, TriPoly considers its compatibility with the sequence reads, as well as the number of recombination events observed by comparing the parental extensions with that of the offspring.

Using quantitative measures, we investigated the quality of haplotype estimates obtained by TriPoly in parent-offspring trios simulated under realistic assumptions with tetraploid *×* tetraploid and tetraploid *×* diploid parents. By comparing our results with the single individual haplotyping methods, we show that substantially better estimates can be obtained by TriPoly for the haplotypes of the progeny, especially at low sequencing depths.

## 2 Method

### 2.1 Specification of a probabilistic model for phasing

In order to establish a probability model for haplotypes, with the sequence reads as data and the base call error and recombination rates as parameters, we must first determine which reads are informative about the phasing. Informative reads need to cover at least two variants, e.g. SNP sites which are heterozygous for at least one of the trio members (*m, f, c*), corresponding to mother, father and the offspring (child). As sites that are homozygous in all trio members retain no phasing information, we discard them from the sequence reads and keep only the base-calls corresponding to the variation positions. Therefore in the first step, the SNP sites, *s* = 1, 2, …, *l*, are detected over a genomic region and the genotypes 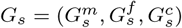 are estimated at these sites, using efficient algorithms such as FreeBayes (Garrison and Marth, 2012). The raw reads of each trio member are then replaced by the so-called *SNP-fragments* of length *l* (Figure 1) that each correspond to a read and contain the numerically coded alleles, i.e. 0, 1, 2 or 3 representing the reference and alternative nucleotides, at the SNP sites covered by that read and ’-’ at positions not called or not covered. To reduce sequencing noise, the positions at which the base-calling quality is lower than a desired threshold can be set to ’-’ as well. Hereafter, by using the term sequence read, *r*, we refer to SNP-fragments that contain at least two determined positions.

**Figure 1:**
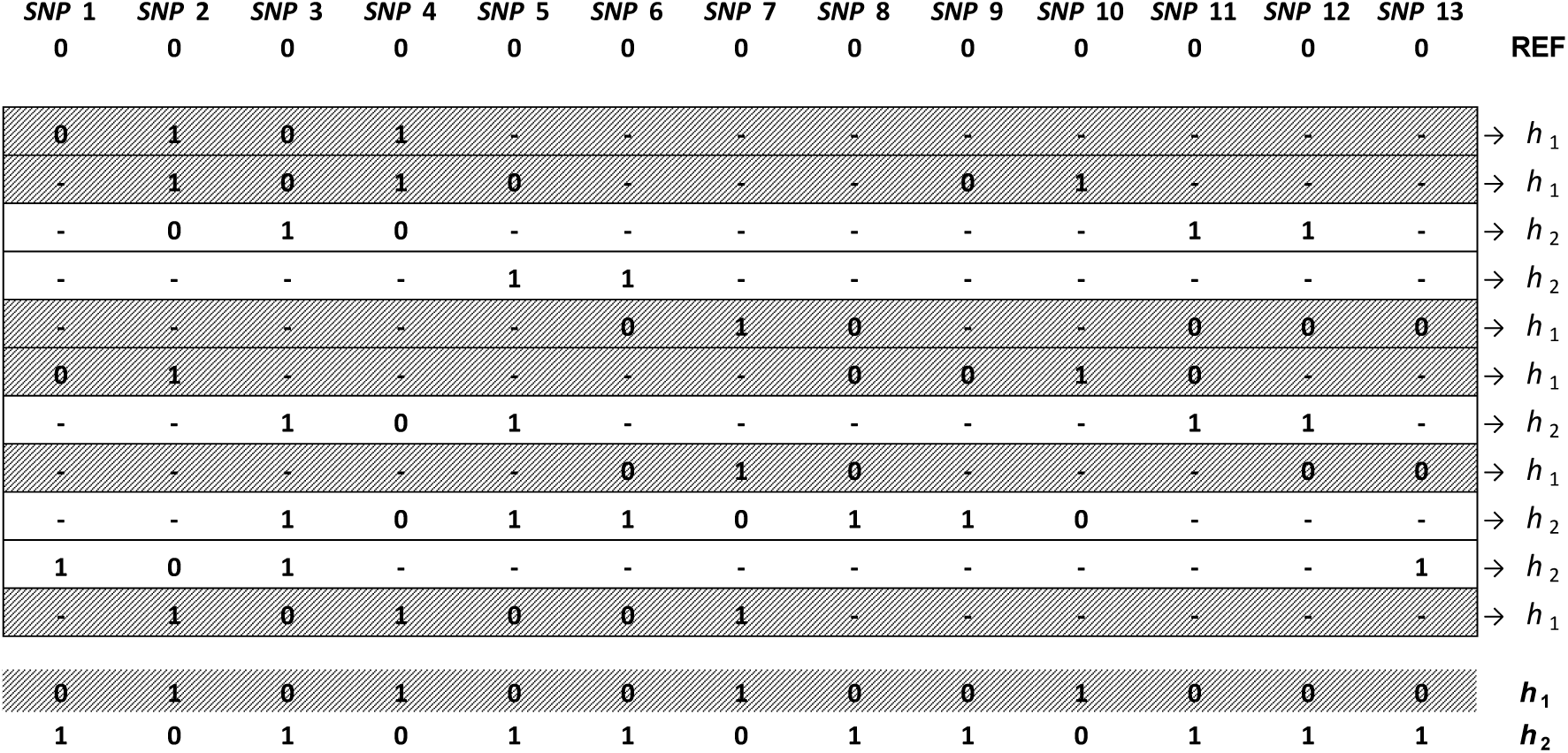
A set of SNP-fragments aligned to a reference and the homologues, *h*_1_ and *h*_2_, from which the fragments are originated. Fragments that have identical variants, specified by 0 (reference) and 1 (alternative), at their overlapping sites are assigned to the same homologue.

In the next step, one should assign the reads to *k* compatible sets in which all of the reads have the same allele at their overlaps, and obtain the consensus sequence of each set to obtain the phasing. As shown in Figure 1, this process is straightforward for diploids in the absence of sequencing errors. In presence of sequencing errors, however, such an assignment of reads to homologues will be possible only if mismatches are allowed. However, allowing mismatches at sites with no error can lead to incorrect haplotype estimates. Polyploidy results in further complexity, as there may be more than one way to assign the reads into *k* > 2 sets even when no error is present. This can happen for instance when several haplotypes are identical in a phasing solution, e.g. in a 3 SNP tetraploid phasing consisting of 4 homologues: 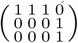 in which three identical 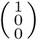 haplotypes are present. In this example, the reads will be compatible with any phasing as long as it contains both 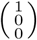 and 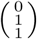 haplotypes regardless of their dosages, e.g. with the phasing 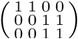. Therefore, probabilistic models must be used to assign the reads to homologues taking into account the uncertainty caused by various phasing possibilities and the presence of errors in the reads.

To account for sequencing errors, we assume an independent binomial error model at each SNP site (Berger *et al.*, 2014) and assign an error vector, 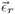, of length *l* to each read containing the probability of erroneous base-calling at the SNP-sites in that read. Using these error probabilities, the probability of maternal, paternal and offspring phasings belonging to a trio, represented by *H*_*m*_, *H*_*f*_ and *H*_*c*_, respectively, can be derived from the set of sequence reads associated with the trio, **R** (consisting of maternal read **R**_*m*_, paternal reads **R**_*f*_ and offspring reads **R**_*c*_). In addition to the reads, we consider meiotic recombination probabilities, *θs*, between SNP *s* − 1 and SNP *s*, represented by vector 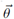, for all *s* > 1 to adjust the probability assigned to each phasing using Mendelian inheritance rules as follows:

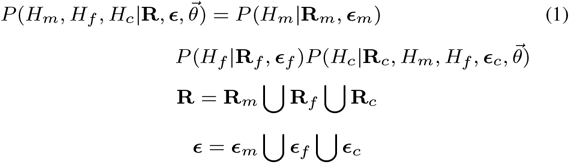

where ***∊***_*m*_, ***∊***_*f*_ and ***∊***_*c*_ are sets of error vectors associated with **R**_*m*_, **R**_*f*_ and **R**_*c*_, respectively. Assuming exchangeability of the offspring, it is straightforward to generalise Equation 1 to include *n* offsprings as:

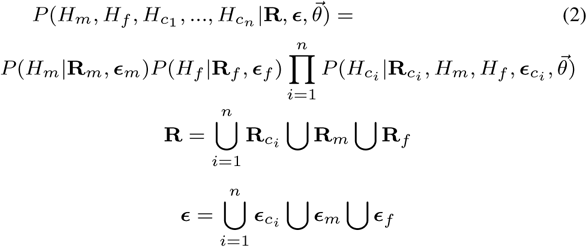

### 2.2 Computational complexity of finding the phasing with the maximum likelihood

By calculating the lefthand side of Equation 1, one can determine the likelihood of each phasing of a trio conditional on its sequence reads. However, as it is instead more convenient to calculate the probability of observing the reads conditional on a phasing (Berger *et al.*, 2014), we use Bayes’ formula for obtaining the phasing likelihoods (Supplementary Methods:Equations 1, 2). To apply this Bayesian approach, one also needs to assign a prior probability to each phasing (Supplementary Methods: Equations 4, 5). The number of recombination events (with some preset recombination rate) can thus be used to assign this prior (Supplementary Methods: Equation 5).

To determine the order of computations needed to find the phasing with the maximum likelihood (determined with the approach described above), we begin by noting that the number of possible phasings of *l* SNPs for a *k*-ploid is bounded in the range:

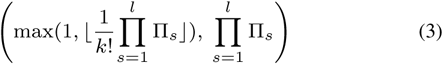

where π_*s*_ denotes the number of possible permutations of the *k* homologues at position *s*. The 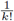 coefficient produces the lower bound, as the numbering of the homologues is arbitrary and therefore each phasing can be obtained by up to *k*! combinations of the single SNP permutations (with *k*! occurring when the phasing is consisted of *k* distinct haplotypes). As an example, for a tetraploid phasing that includes 3 SNPs (1 *≤ s ≤* 3) with genotypes: *G*_1_ = (1, 1, 0, 1), *G*_2_ = (0, 0, 1, 0) and *G*_3_ = (0, 0, 1, 0), Equation 3 gives lower and upper bounds equal to 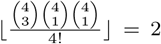 and 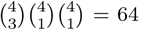, respectively, while 5 distinct phasings: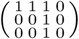, 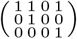, 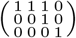 and 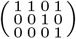, 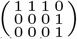 are actually possible, yielded by 12, 24, 12, 12 and 4 combinations of the single SNP permutations, respectively.

With parental ploidy levels *k*_*m*_, *k*_*f*_ and parental sequencing depths *c*_*m*_, *c*_*f*_, calculating the probability of each parental phasing conditional on its reads requires *O*(*k_p_lc_p_*) computations for *p ∈ {m, f}*, as each determined allele in the reads must be compared to the corresponding allele on each of the *k_p_* homologues and each SNP has been on average called in *C*_*p*_ reads. Assuming no recombination, at most 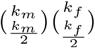 distinct haplotypes can be passed from the parents to the offspring through balanced meioses, yielding an offspring ploidy level *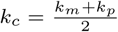* Similar to the parental phasings, each offspring phasing requires *O*(*k*_*c*_*lc*_*c*_) computations to calculate its probability conditional on the offspring reads at an average depth of *cc*. Therefore, from Equation 3 it follows that a total computational cost of *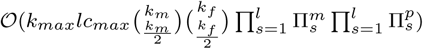* is required to calculate Equation 1 assuming no recombination, with *kmax* the maximum parental ploidy level, i.e. max(*k*_*m*_, *k*_*f*_), and *c*_*max*_ the maximum sequencing depth for the trio members.

Allowing for recombination, different homologues may be passed to the offspring at each SNP position. To take all possible transmissions into account, we have to enumerate them separately at each SNP position. Thus, the order of computations increases to 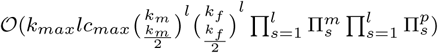.

### 2.3 TriPoly algorithm

At fixed ploidy levels, the computational cost of the brute-force approach calculated in Section 2.2 grows linearly with sequencing depth, but exponentially with the number of SNPs *l*, rapidly rendering this approach intractable.

To overcome this problem, we perform SNP-by-SNP reconstruction of haplotypes, starting from the leftmost SNP in the target region and keeping only a few most likely phasing extensions to the next SNP at each step (Figure 2). Following this approach, one will end up with a limited number of phasings that have passed the selection criteria during the extension procedure from *s* = 1 to *s* = *l*. Assuming the selection procedure effectively keeps the number of accepted solutions at each extension step bounded above by *E*_*m*_ and *E*_*f*_ for the mother and the father, respectively, the number of trio phasings at each extension will be bounded above by 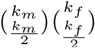 and the total complexity will decrease to 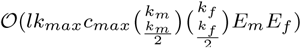. This greedy method is therefore linear in terms of the number of SNPs, *l*. With parental ploidy levels, *k_p_* for *p∈ {m, f}*, in the range of 2 to 12 (covering most of the naturally occurring cases of polyploidy), 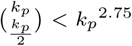 and therefore this cost grows with a rate of *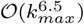* with ploidy level.

**Figure 2:**
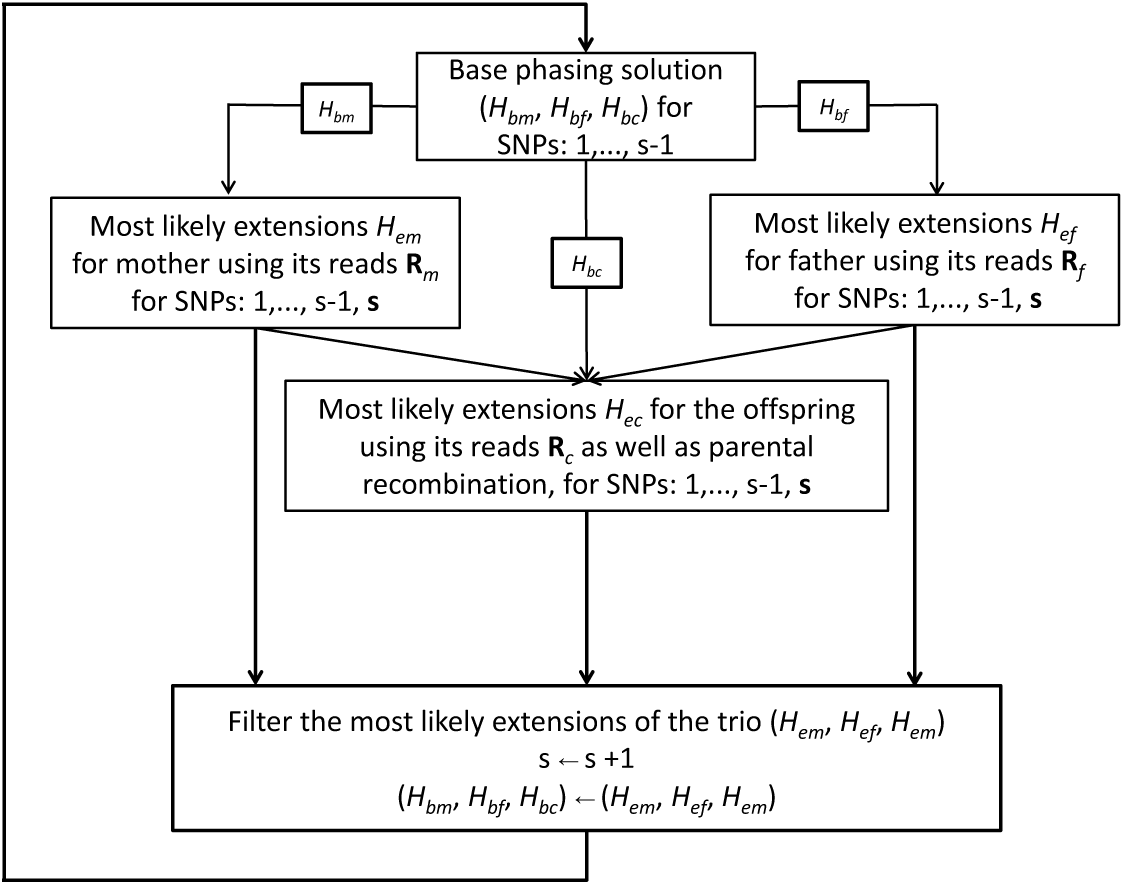
Overview of the SNP by SNP haplotyping method implemented in TriPoly for a trio consisted of two parents and one offspring.

To implement this approach, which we call *TriPoly*, we employ the *branching* and *pruning* steps explained in HapTree algorithm (Berger *et al.*, 2014), as shown in Figure 3. Starting at SNP site *s* = 1, its alleles for each parent and for the offspring are used as the base parental and offspring phasings, *H*_*bp*_ and *H*_*bc*_. The phasing is then extended step by step from SNP *s* – 1 to SNP *s* for *s ≥* 2, until all SNPs have been phased according to the algorithm outlined in Supplementary Methods: Algorithm 1. At each extension step, branching and pruning (Supplementary Methods: Procedure 3 and Supplementary Methods: Procedure 4) allow the algorithm to work with a limited number of phasing solutions. This approach can be easily extended to include several offspring at the same time using Equation 2, a detailed description of which is given in Supplementary Methods, A.

**Figure 3:**
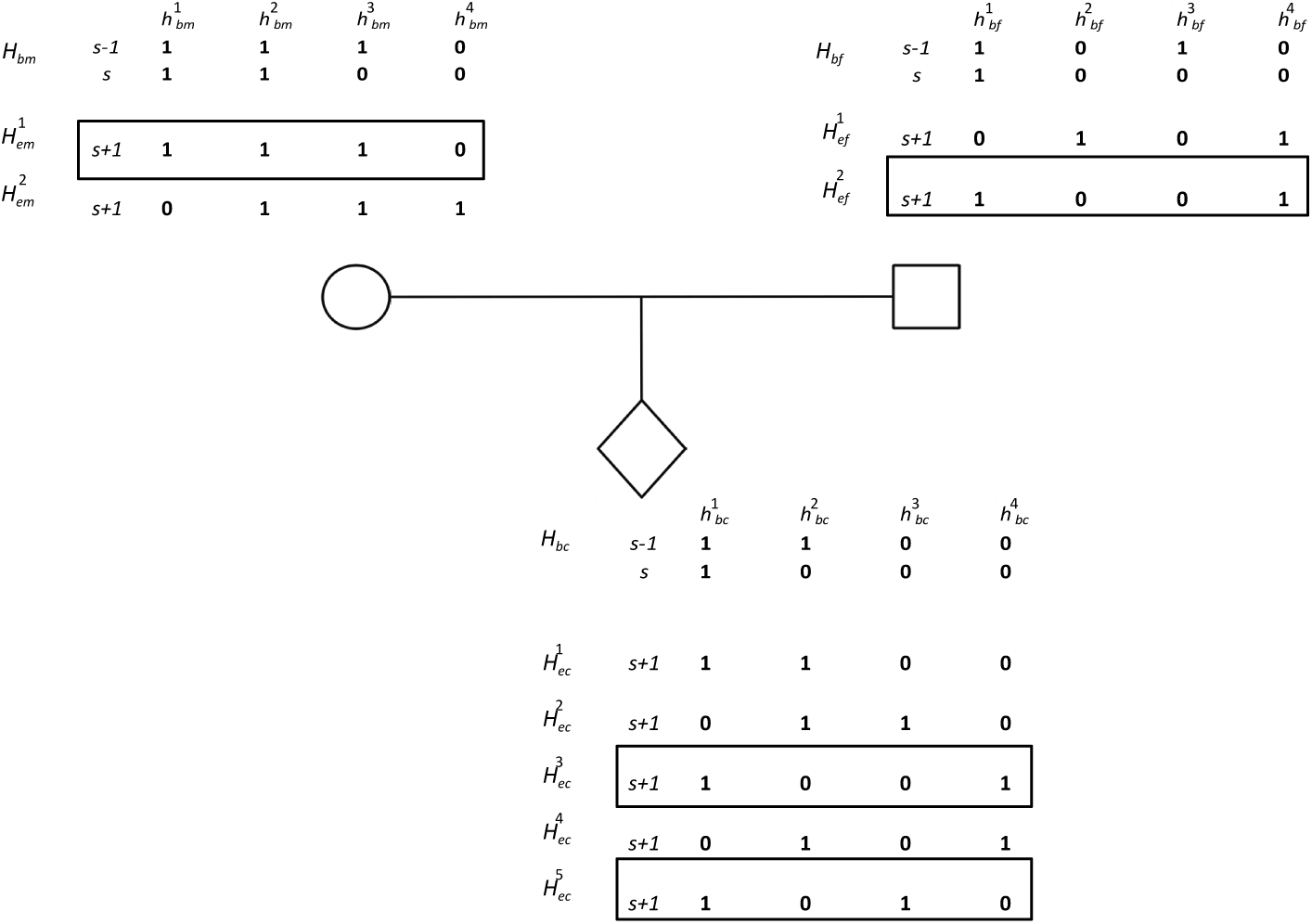
An example of the branching step for a trio: mother and father base phasings, *H*_*bm*_ and *H*_*bf*_, ending at SNP *s* – 1 are extended to SNP *s* using sequence reads. Extensions *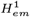* and *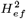* that have a posterior probability larger than the branching threshold, *ρ*, are used to extend the offspring base phasing *H*_*bc*_ by transmitting their alleles at *s*, assuming the offspring homologues 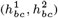 have maternal origin and *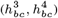* are of paternal origin. Thus,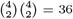 transmissions are possible from which those not compatible with the offspring genotype at *s* are discarded. Also in case several transmissions result in the same phasing for the offspring, only one transmission is considered that implies the minimum number of recombinations. Using the offspring reads as data and the recombination probability as prior, a Bayesian probability is assigned to each offspring extension. Extensions *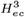* and *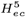* that pass *ρ* yield candidate trios extensions:*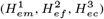* and 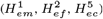. The probability of each trio extension is determined using Equation 1 and is compared against *ρ* to choose the final extensions.

Note that this approach assumes working on the so-called phasing *blocks*, i.e. genomic regions in which each SNP, *s*, is connected to at least one other SNP, *s*^*'*^, through at least one of the reads in **R**. In case the sequencing reads do not satisfy this condition for the whole set of SNPs in the region, it is straightforward to divide the SNP set into blocks and phase each block separately, with the phasing being interrupted between the blocks.

## 3 Experimental setup

### 3.1 Simulation of polyploid trios

We evaluated the performance of TriPoly, as well as three state-of-theart single individual haplotyping algorithms: HapCompass, SDhaP and HapTree, using synthetically generated sequence data for parent-offspring trios. To this end, maternal and paternal genomes were independently simulated from a common reference using *Haplogenerator* (Motazedi *et al.*, 2017), and offspring genomes were generated by passing recombinant parental chromatids at random considering a Poisson stochastic model for meiosis (see Supplementary Methods, B for the details). In our simulations, we set the recombination rate (*λ* in Supplementary Methods: Equation 7) to 3.07 *cM/Mb*, corresponding to the average recombination rate in potato (Bourke *et al.*, 2015; Felcher *et al.*, 2012). Using this approach, genomic regions of length 10 kb where simulated for 100 independent trios of tetraploid (*k*_*m*_ = *k*_*f*_ = *k*_*c*_ = 4) potato (*Solanum tuberosum*, 2*n* = 4*x* = 48), based on 100 regions randomly selected from PGSC-DM genome, chromosome 5 (release version 4.03) (Genome Sequencing Consortium *et al.*, 2011) using a lognormal model to simulate genomic variation (Motazedi *et al.*, 2017). To fit the lognormal model, the SNP density of each parent was determined from empirical data (Uitdewilligen *et al.*, 2013) as described in (Motazedi *et al.*, 2017), resulting in a mean distance of 21 bp between neighbour SNPs with a standard deviation of 27 bp. The proportion of each parental marker type: simplex, duplex, triplex and quadruplex, in the total set of markers was also determined from (Uitdewilligen *et al.*, 2013) to be 0.5, 0.23, 0.14 and 0.13, respectively.

We also simulated crosses of diploid (2*n* = 2*x* = 22) and tetraploid (2*n* = 4*x* = 44) banana (*Musa acuminata*), with the female parent being the tetraploid as the pollen of tetraploid banana is hardy viable (Fortescue and Turner, 2004). In practice, commercial triploid bananas (*kc* = 3) are produced by such hybridisations (*k*_*m*_ = 4, *k*_*f*_ = 2), which have high consumer preference as their parthenocarpic fruits lack the large, hard seeds of fertilisation-induced fruits of diploid and tetraploid sorts.

We used the sequence of chromosome 10 from the reference genome of DH-Pahang (a double-haploid *M. acuminata*) (D’Hont *et al.*, 2012), release version 2 (Martin *et al.*, 2016), to simulate banana trios, applying the lognormal model to generate SNPs. To fit the model, we set the average SNP frequency to 1 per 200 bp with a standard deviation of 1194 bp, so that we do not get many uninformative reads (Section 2.1) while the predicted average distance of 1394 bp between DH-Pahang SNPs (Droc *et al.*, 2013) lies one standard deviation away from the considered average. As 1% recombination rate has been reported to correspond to 100 to 400 kb physical distance for banana (except at regions close to the centromere) (Pillay *et al.*, 2012, p. 130), we applied an average recombination rate of 0.04 *cM/Mb* simulating meiosis. The proportions of parental marker types were set the same as that of potato.

For each simulated individual, simulation of the sequence data and variant calling was performed using conventional tools, explained in detail in Supplementary Methods, C.

### 3.2 Measures of Phasing Estimation Quality

Knowing the true haplotypes in simulations, one can evaluate the performance of haplotyping methods by using measures that directly compare the estimates to the true haplotypes. We used the *reconstruction rate* (*RR*) (Geraci, 2010) and the *pair-wise phasing accuracy rate* (*PAR*) (Browning and Browning, 2011) to evaluate the accuracy, and the SNP missing rate (SMR) (Motazedi *et al.*, 2017) as well as the number of gaps per SNP (NGPS) to evaluate the completeness and continuity of haplotyping.

The first measure, RR, has been defined for diploids as the proportion of correctly phased markers in the phasing estimate of the target region (Geraci, 2010). However, to apply it for polyploids we have to generalise its mathematical formulation as haplotypes are not necessarily complementary in polyploids, making multiple correspondences possible between the original and estimated haplotypes.

Let 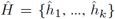 be the estimated phasing and *H* = *{h*_1_, …, *h*_*k*_*}* be the correct phasing of a region containing *l* SNPs. We define RR as:

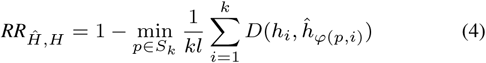

where *S*_*k*_ represents the permutation group on *{*1, …, *k}* and *ϕ* denotes the group action on *{*1, *…, k}*. In this definition, *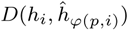* is the Hamming distance defined as:

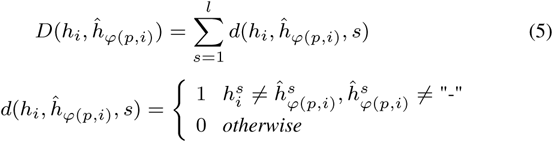

where 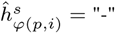 means that SNP *s* has not been phased in 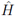.

As an alternative measure of estimation accuracy, PAR is defined as the proportion of all SNP pairs for which the inferred phasing is correct. While RR is an overall measure of the accuracy of local phasing, i.e. the phasing inferred from the estimated haplotypes for a few adjacent SNPs along the target region, PAR primarily shows the accuracy of long range phasing as it is highly affected by chimeric elongations of the haplotypes during estimation, i.e. the elongation of a homologue by part of another homologue.

TriPoly: haplotype estimation for polyploids using sequence data of related individuals 5

As haplotyping methods sometimes report phasings with high SNP exclusion, which nevertheless can have high RR and PAR, the average proportion of SNPs left out in the phasing estimates of each method was calculated as SMR to show the method’s phasing completeness. Besides, in order to show how much fragmented the phasing estimates are for each method, which phenomenon is not reflected in RR, PAR or SMR, the average number of interruptions, i.e. the number of blocks minus one, in the estimates of each method was calculated and normalised by the number of SNPs, *l*, as NGPS. Defined in this way, NGPS measures the continuity of phasing for each method. All of the calculations to obtain these quality measures were performed using *hapcompare* (Motazedi *et al.*, 2017).

To quantify the effect of haplotyping method on the quality measures in each simulated population, accounting for the effect of sequencing depth and random variation among the simulated families, we built regression models for each measure including the estimation method as predictor.

## 4 Results

We used simulated genomes and sequence reads to assess the performance of TriPoly, HapCompass, HapTree and SDhaP in trios of tetraploid potato and tetraploid-diploid-triploid banana. To quantify the assessment, we used pairwise phasing accuracy rate (PAR) and reconstruction rate (RR) (Section 3.2) as measures of log-range and local phasing precision, respectively, and the number of gaps per SNP (NGPS) in each estimate as measure of phasing continuity. The fraction of unphased SNPs was also reported as SNP missing rate (SMR) for each haplotyping method to show the method’s phasing completeness. For each simulation scenario, we built a regression model to investigate the dependency of each of these measures on the haplotyping method and sequencing depth.

All of the analyses were run using 2.90 GHz Intel Xeon processors. A time-limit of 1500 seconds was set for each haplotyping method during simulations, not to consume too much of the shared computational resources in case estimation became prohibitively difficult (Motazedi *et al.*, 2017). To achieve time-memory efficiency, we set the branching threshold of TriPoly, *ρ*, to 0.2 and its pruning threshold, *κ*, to 0.94. Besides, we forced TriPoly to keep no more than 11% of all possible phasing extensions at each step in case the pruning had not been able to discard as many with the value chosen for *κ*.

### TriPoly improves the quality of phasing between neighbouring SNPs for the offspring

The results of regression analysis showed 11% and 24% increases in RR by using TriPoly compared to the other methods for the banana and potato offspring, respectively (Supplementary Tables S1-S2). These observed improvements in local phasing show that parental transmission is informative even for phasing between nearby SNPs, in which case the SNPs can be contained within a single read and therefore the reads dominate the phasing likelihood. This information is especially advantageous when the offspring is sequenced at low depth (Figure 4). TriPoly did not increase RR for the parents (Supplementary Figures S2- S3).

**Figure 4:**
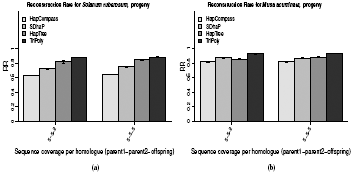
Average reconstruction rates (RR) for the progeny in the 100 trios simulated for a) potato and b) banana, obtained by HapCompass, SDhaP, HapTree and TriPoly at various sequencing depths.

Among the single individual haplotyping methods, HapTree yielded the closest RR, but its accuracy was more variable at low depth compared to the other methods (Figure 4).

### TriPoly increases markedly the accuracy of phasing between distant SNPs for the offspring

The regression analysis of PAR showed that at the same SMR, the fraction of correct phasings between distant SNPs is around 33% and 42% higher in the TriPoly estimates for banana and potato offspring, respectively, compared to the other methods (Supplementary Tables S1-S2, Supplementary Figure S1). Besides, this increase was more manifest at low sequencing depths (Figure 5). TriPoly was not able to increase PAR for the parents (Supplementary Figures S2- S3).

**Figure 5:**
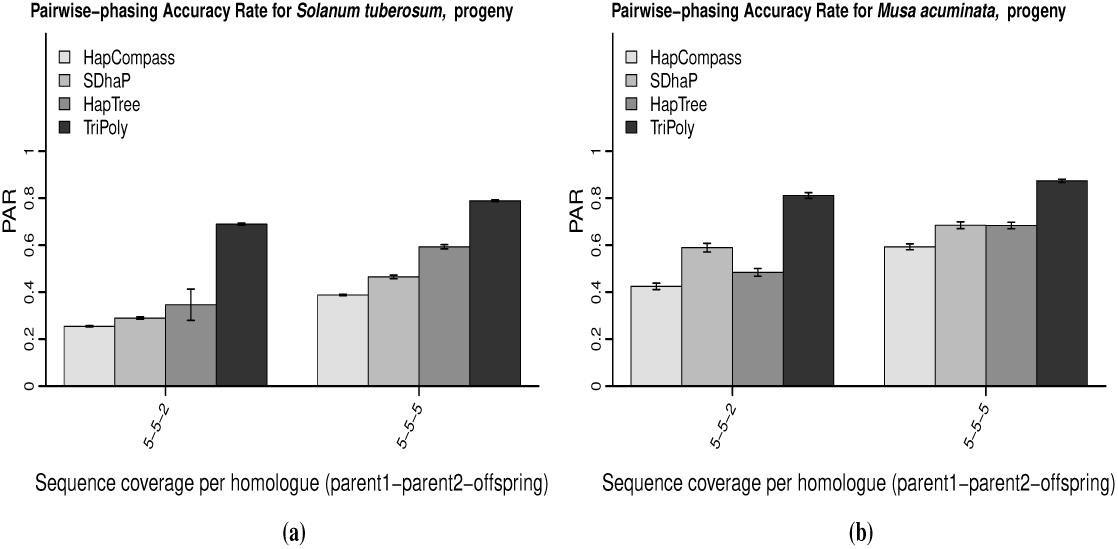
Average pairwise-phasing accuracy rates (PAR) for the progeny in the 100 trios simulated for a) potato and b) banana, obtained by HapCompass, SDhaP, HapTree and TriPoly at various sequencing depths.

While sequence reads contain phasing information for the SNPs contained in them, this information hardly goes beyond nearby SNPs with short reads that span just a few hundred bases. The accuracy of phasing is therefore gradually decreased as the phasing is extended to include more distant SNPs in a block, due to the fact that chimeric extensions become more likely with spurious overlaps between the erroneous reads. By penalising recombination events through the considered small recombination probability (Supplementary Methods: Equation 6), TriPoly tends to reduce the chance of chimeric extensions and markedly improves the precision of phasing between distant SNPs in the offspring.

Similar to RR, the PAR scores closest to TriPoly were obtained by HapTree, but HapTree results were more variable in accuracy at low depths.

### Fewer phasing interruptions are introduced in the haplotypes estimated by TriPoly

As explained in Section 3.2, in single individual haplotyping the phasing is interrupted between two SNPs if there is no read that connects the two by covering both. However, when the reads do not contain enough phasing information for some SNPs, parental transmission can be still informative to prefer one phasing extension to another (Supplementary Methods, A) resulting in less phasing interruptions. The regression analysis of NGPS showed that the haplotypes obtained by TriPoly were significantly less interrupted compared to the other approaches, notably for banana (Supplementary Tables S1-S2, Figure 6).

**Figure 6:**
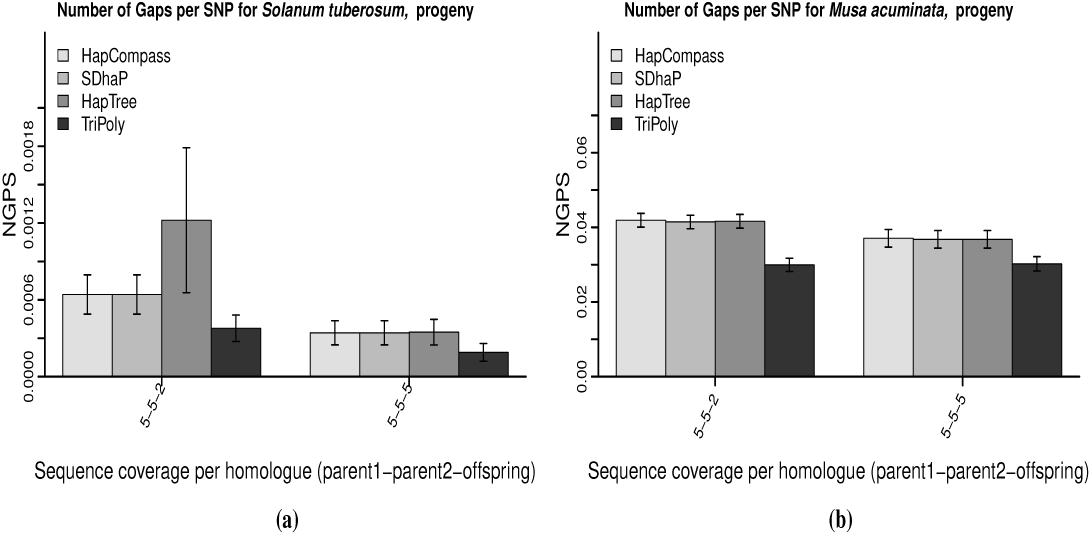
Number of Gaps per SNP (NGPS) in the phasing estimates of the progeny from the 100 trios simulated for a) potato and b) banana, using HapCompass, SDhaP, HapTree and TriPoly at various sequencing depths.

At lower SNP densities, the average distance between subsequent SNPs will be larger and this can increase the number of reads uninformative for phasing (Section 2.1). As a results, more interruptions can be introduced in the haplotypes reconstructed from short reads (Motazedi *et al.*, 2017). TriPoly proves to be beneficial in such low SNP density situations, reflected in the notable decrease in NGPS for banana compared to the slight decrease for potato which has a high SNP density.

Finally, the high standard deviation of NGPS for HapTree stands out in Figure 6-a, which is a reflection of its high failure rate at low sequencing coverages for tetraploid potato (Motazedi *et al.*, 2017). As all of the SNPs belonging to a failed block are excluded from the final phasing, the NGPS will be more varying across the simulated trios due to chance failures.

### TriPoly has the smallest memory consumption and finishes estimation during a time comparable to that required by the other methods

As processing large genomic regions usually requires considerable amounts of CPU time and memory, it is important for a haplotyping algorithm to be efficient in terms of these two resources. Therefore, we measured the computation time and memory consumption of TriPoly for the simulated potato and banana trios at the applied sequencing depths and compared it to those of HapCompass, HapTree and SDhaP. As shown in Figure 7, TriPoly is the most memory-efficient algorithm compared to the others, while it requires more time compared to HapCompass and SDhaP for potato. However, the amount of time required by TriPoly was still not very far from that needed by the other algorithms.

**Figure 7:**
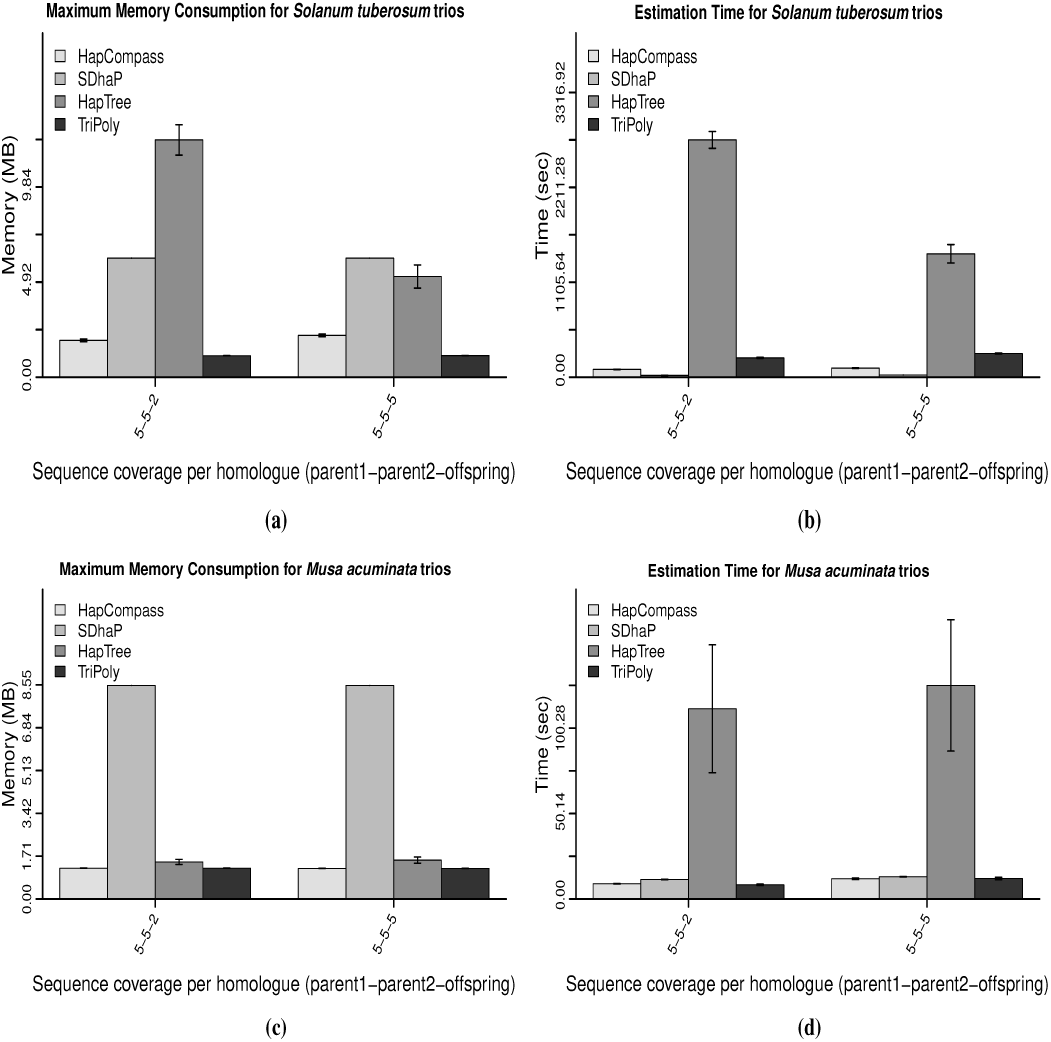
Memory consumption of the haplotyping algorithms (a, c), and the running time of each (b, d) for the 100 simulated banana and potato trios, respectively, at various sequencing depths.

## Conclusion and Discussion

We propose a novel approach, called TriPoly, for estimating haplotypes in polyploid parent-offspring trios using NGS data while taking haplotype transmission from the parents to the progeny into account. TriPoly reconstructs the phasing of the SNPs over a genomic region simultaneously for the parents and for the offspring setting out with the SNP site that has the smallest coordinate in the region, adding one SNP to the phasing at each step and greedily selecting the most likely extensions for the next extension step conditional on the sequence reads and recombination events. Through realistic simulations, we show that TriPoly significantly improves the haplotyping accuracy for the offspring by 11-42% compared to single individual approaches: HapCompass, SDhaP and HapTree. Besides, we show that TriPoly estimates are more continuous compared to the other methods when the SNP-density is low. TriPoly in also an efficient algorithm in terms of the memory consumption and CPU-time.

In contrast to HapCompass, SDhaP and HapTree, TriPoly provides an option to include all of the SNPs, i.e. including those homozygous or missing for an individual, in the output. In this way, the haplotypes can be compared in an F1-population and the segregation patterns can be easily investigated. Moreover, haplotypes reported in this format can be coded as multi-allelic markers to be used in genetic analyses. Besides, TriPoly accepts input in the more convenient format of multi-sample BAM and VCF files, compared to the other methods that either require one-sample BAM/VCF (HapCompass) or the SNP-fragment matrix in place of the mapped reads (SDhaP and HapTree).

While TriPoly increases the accuracy of phasing for the offspring in a trio by incorporating parental recombination probabilities in the phasing likelihood (Equation 1), it assumes exchangeability of the offspring in families with more than one offspring (Equation 2), which ignores the phasing information conveyed by an offspring about the others. By implementing more complex joint likelihood models, we can expect to see an enhancement in haplotyping accuracy for larger families, both in the parental haplotypes as well as in the progeny haplotypes. However, the computational burden is definitely a challenge in implementing such an approach. Another potential improvement in TriPoly is the phasing of the parents, the accuracy of which was shown to be inferior to that obtained by HapTree. An iterative approach of keeping a few surviving TriPoly solutions for the whole target region as the starting point for an Expectation Maximisation (EM) routine can be a way to tackle this problem, resulting in a refined set of most likely haplotypes in the population to which the reads of each individual can be mapped back to find its specific phasing. Like the joint likelihood approach, the computational challenge will be an important consideration here.

## Acknowledgement

The authors thank the graduate school Experimental Plant Sciences (EPS), Wageningen University and Research (WUR) for funding this work and Peter Bourke (WUR-Plant Breeding) for his valuable comments.

## Software

TriPoly has been implemented in Python 3.5.2 (also compatible with Python 2.7.3 and higher) and can be freely downloaded at *www.bif.wur.nl*.

---

Supplementary Figure S1: Suppfig1.pdf: Average SNP Missing Rates (SMR) in the phasing estimates of the progeny from the 100 trios simulated for a) potato and b) banana, using HapCompass, SDhaP, HapTree and TriPoly at various sequencing depths.

Supplementary Figure S2: Suppfig2.pdf: Plots of RR, PAR, NGPS and SMR obtained by HapCompass, SDhaP, HapTree and TriPoly for the parents in the 100 simulated *M. acuminata* trios.

Supplementary Figure S3: Suppfig3.pdf: Plots of RR, PAR, NGPS and SMR obtained by HapCompass, SDhaP, HapTree and TriPoly for the parents in the 100 simulated *S. tuberosum* trios.

**Table S1:** 95% Confidence intervals for regression of quality measures on haplotype estimation variables for *M. acuminata*

|  | RR | PAR | SMR | NGPS |
| --- | --- | --- | --- | --- |
| Intercept | 0.813(0.803;0.823) | 0.443(0.42;0.466) | 0.624(0.613;0.636) | 0.0412(0.0377;0.0448) |
| COV 5-5-5 | 0.008(0;0.015) | 0.131(0.113;0.15) | -0.256(-0.263;-0.249) | -0.0035(-0.0057;-0.0013) |
| SDhaP | 0.052(0.041;0.063) | 0.128(0.102;0.155) | 0.002(-0.008;0.011) | -0.0004(-0.0035;0.0027) |
| HapTree | 0.052(0.041;0.062) | 0.075(0.049;0.101) | 0.001(-0.008;0.011) | -0.0003(-0.0034;0.0028) |
| TriPoly | 0.113(0.102;0.123) | 0.334(0.308;0.36) | 0.012(0.003;0.022) | -0.0094(-0.0125;-0.0063) |

**Table S2:**
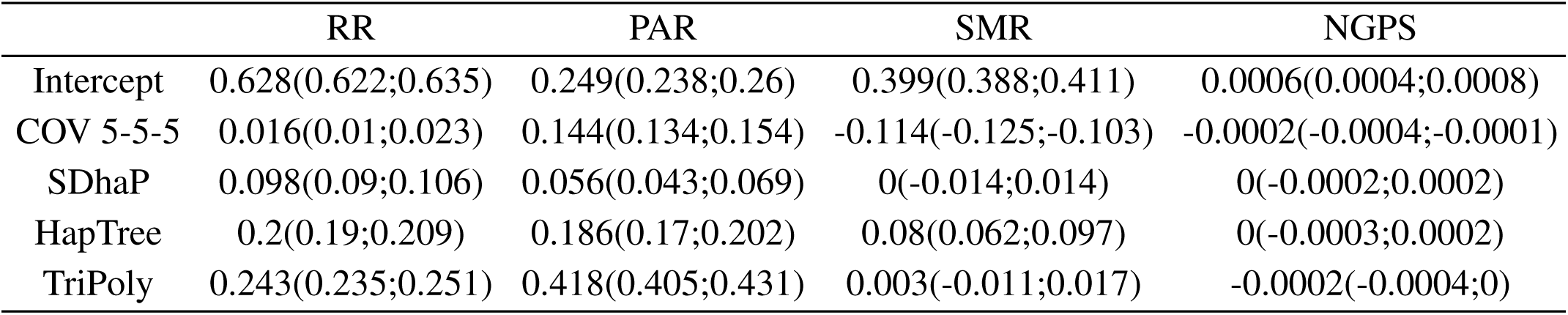
95% Confidence intervals for regression of quality measures on haplotype estimation variables for *S. tuberosum*

|  | RR | PAR | SMR | NGPS |
| --- | --- | --- | --- | --- |
| Intercept | 0.628(0.622;0.635) | 0.249(0.238;0.26) | 0.399(0.388;0.411) | 0.0006(0.0004;0.0008) |
| COV 5-5-5 | 0.016(0.01;0.023) | 0.144(0.134;0.154) | -0.114(-0.125;-0.103) | -0.0002(-0.0004;-0.0001) |
| SDhaP | 0.098(0.09;0.106) | 0.056(0.043;0.069) | 0(-0.014;0.014) | 0(-0.0002;0.0002) |
| HapTree | 0.2(0.19;0.209) | 0.186(0.17;0.202) | 0.08(0.062;0.097) | 0(-0.0003;0.0002) |
| TriPoly | 0.243(0.235;0.251) | 0.418(0.405;0.431) | 0.003(-0.011;0.017) | -0.0002(-0.0004;0) |

